# Spinal cholinergic interneurons differentially control motoneuron excitability and alter the locomotor network operational range

**DOI:** 10.1101/135723

**Authors:** Maria Bertuzzi, Konstantinos Ampatzis

## Abstract

While cholinergic neuromodulation is important for locomotor circuit operation, the specific neuronal mechanisms that acetylcholine employs to regulate and fine-tune the speed of locomotion are largely unknown. Here, we show that cholinergic interneurons are present in the zebrafish spinal cord and differentially control the excitability of distinct classes of motoneurons (slow, intermediate and fast) in a muscarinic dependent manner. Moreover, we reveal that m2-type muscarinic acetylcholine receptors (mAChRs) are present in fast and intermediate motoneurons, but not in the slow motoneurons, and that their activation decreases neuronal firing. We also provide evidence that this configuration of motoneuron muscarinic receptors serves as the main intrinsic plasticity mechanism to alter the operational range of motoneuron modules. These unexpected findings provide new insights into the functional flexibility of motoneurons and how they execute locomotion at different speeds.

## Introduction

Neural networks in the spinal cord are responsible for the generation and execution of movements (Goulding, 2009; Grillner, 2003; Grillner and Jessell, 2009; Kiehn, 2006). These spinal cord networks are organized in distinct functional microcircuit modules (Ampatzis et al., 2014; Bagnall and McLean, 2014) with defined operational ranges, whose sequential activation increases the speed of locomotion (Ampatzis et al., 2014). The activity of the spinal neuronal microcircuits is regulated by a range of neuromodulatory systems (Miles and Sillar, 2011) that permit motoneurons to adjust their final motor output. One such prominent neuromodulatory system is the cholinergic system (Miles et al., 2007; Zagoraiou et al., 2009). In rodents, cholinergic neuromodulation increases motoneuron excitability (Miles et al., 2007; Deardorff et al., 2014) in a task-dependent manner (Zagoraiou et al., 2009), which is exclusively mediated by cholinergic V0c interneurons (Zagoraiou et al., 2009). Whether cholinergic interneurons are present in the zebrafish spinal cord, and how they regulate the activity of the distinct motoneuron modules (slow, intermediate and fast) during locomotion is, however, unknown.

In mammals, acetylcholine release increases motoneuron excitability (Chevallier et al., 2006; Hornby et al., 2002; Ireland et al., 2012; Miles et al., 2007). Although m2 type metabotropic muscarinic acetylcholine receptor (m2-mAChR) activation is reported to mediate mammalian motoneuron hyperexcitability (Deardorff et al., 2014; Miles et al., 2007; Zagoraiou et al., 2009), numerous studies also demonstrate m2-mAChR-mediated inhibitory actions, in several neuronal populations (Brown, 2010; Felder, 1995; Hosey, 1992), including motoneurons (Kurihara et al., 1993). Moreover, m2-mAChRs are found to be predominantly expressed in large motoneurons (Welton et al., 1999). This suggests that only a subset of motoneurons are sensitive to cholinergic modulation via the m2-mAChRs. In order to resolve the question of whether m2 receptors are present in all motoneuron pools (slow, intermediate and fast), and determine how activation of the m2-mAChRs influences the motoneuron functionality, we used the accessible neuro-muscular configuration of the adult zebrafish (Ampatzis et al., 2013).

Using a combination of anatomical, electrophysiological, pharmacological, *ex vivo* and *in vivo* behavioral approaches in adult zebrafish, we uncover a previously unidentified population of spinal cord cholinergic interneurons that is analogous to mammalian V0c interneurons. Furthermore, we provide the first functional evidence that cholinergic modulation acts as a plasticity mechanism to differentially regulate the excitability and operational range of separate motoneuron modules through activation of distinct mAChR subtypes.

## Results

### Zebrafish spinal cholinergic system organization

We first examined the distribution pattern of cholinergic terminals (VAChT^+^) in the adult zebrafish spinal cord. Multiple cholinergic terminals were detected in the dorsal horn, neuropil and motor column (Figure 1A). Using the accessible neuro-muscular configuration of the adult zebrafish, we dissected distinct functional motoneuron pools (Ampatzis et al., 2013). All retrogradely traced secondary motoneuron types (slow, intermediate and fast) received abundant cholinergic innervation (Figure 1B). To determine whether this innervation differs among functionally different pools of secondary motoneurons, we analyzed the number of cholinergic terminals (VAChT^+^) in close proximity to motoneuron cell bodies (see Materials and Methods). Fast motoneurons receive a significantly higher number of putative cholinergic inputs than the intermediate and slow motoneurons (One-way ANOVA, F_2,64_ = 16.69, p < 0.0001, n = 67 neurons; Figure 1C). Furthermore, our analysis showed that these cholinergic synapses on motoneurons do not display the typical morphology of the mammalian “c-boutons” (Zagoraiou et al., 2009; Herron and Miles, 2012; Skup et al., 2012; Deardorff et al., 2014). To investigate the origin of this input, we retrogradely labeled the brain neurons descending to spinal cord (fluorescent Dextran tracer; see Material and Methods) and subsequently immunolabeled cholinergic neurons (Choline Acetyltransferase (ChAT) immunoreactivity; n = 8 brains). Our analysis revealed that none of these neurons were cholinergic as shown by the co-localization experiments (Supplementary file 1A-H). Moreover, few retrogradely labeled neurons in the initial part of the spinal cord were found to be cholinergic (ChAT^+^) (Supplementary file 1I).

**Figure 1.**
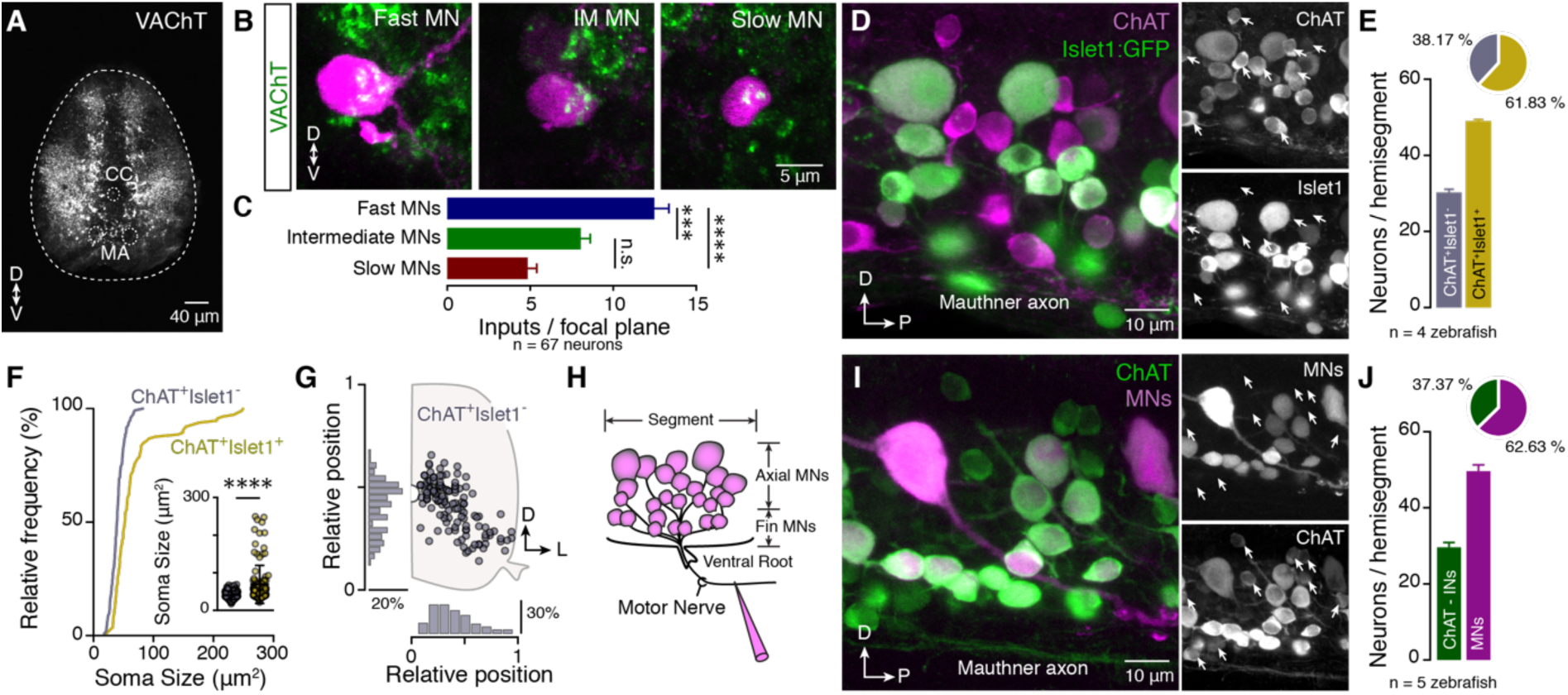
Organization of spinal cholinergic system. (**A**) Distribution of cholinergic terminals within the adult zebrafish spinal cord. (**B**) Single optical sections show the cholinergic synapses onto different types of motoneurons. (**C**) Cholinergic (ACh) input is different between motoneuron (MN) classes, (n_slow_ = 10, n_interm_ = 27, n_fast_ = 30 neurons; n = 5 zebrafish). (**D**) Whole mount spinal cord image shows that cholinergic interneurons ChAT^+^Islet1^-^ exist in adult zebrafish. Arrows indicate the ChAT^+^Islet1^-^ neurons (Cholinergic interneurons). (**E**) Quantification of the adult zebrafish cholinergic neurons, in *Islet1:GFP* line, per spinal cord hemisegment (n = 4 zebrafish). (**F**) Cholinergic interneurons (ChAT^+^Islet1^-^) have smaller soma sizes compared to motoneurons. (**G**) Distribution pattern of cholinergic interneurons (ChAT^+^Islet1^-^) within the adult zebrafish spinal cord. (**H**) Drawing showing the methodological approach to backfill all the motoneurons (axial and fin) exiting the spinal cord through the ventral root by injecting a retrograde tracer (Dextran). (**I**) Whole mount spinal cord immunohistochemistry reveals that a fraction of ChAT^+^ neurons are not backfilled motoneurons. Arrows indicate ChAT^+^MN^-^ neurons (Cholinergic interneurons; n = 5 zebrafish). (**J**) Quantification of the adult zebrafish cholinergic neurons, per spinal cord hemisegment (n = 5 zebrafish), reveals similar number of cholinergic interneurons as in the *Islet1:GFP* line. CC, central canal; D, dorsal; L, lateral; MA, Mauthner axon; P, posterior; V, ventral. Data are presented as mean ± SEM; ***p < 0.001; ****p < 0.0001; n.s., non-significant.

Having established that spinal cord cholinergic innervation is not provided by brain descending neurons, we hypothesized that spinal cord cholinergic interneurons account for the strong cholinergic input to motoneurons. This hypothesis is supported by the morphology of zebrafish motoneurons, which lack axonal collaterals (Ampatzis et al., 2013), that are recurrent axons that branch off from the main axon. In mammals, motoneuron collaterals form synaptic contacts with the Renshaw cells and other motoneurons (Eccles et al., 1954; Cullheim et al., 1977). Renshaw cells have not been reported in the zebrafish spinal circuits, and zebrafish motoneuron-motoneuron communication is meditated through dendro-dendritic electrical gap junctions (Song et al., 2016). We tested our hypothesis using two complementary approaches. First, we employed the *Islet1:GFP* zebrafish line, in which all motoneurons express green fluorescent protein (GFP; Uemura et al., 2005) and labeled both motoneurons and cholinergic interneurons through ChAT immunoreactivity (Figure 1D). We identified a significant fraction (38.17 ± 2.523%, Figure 1E), of small to medium size non-motoneuron cholinergic neurons (ChAT^+^Islet1^-^, 40.13 ± 1.038 μm^2^; Figure 1F), which were distributed throughout the motor column (Figure 1G). In comparison, identified motoneurons were larger (ChAT^+^Islet1^+^, 69.04 ± 4.729 μm^2^; Figure 1F). To further confirm that all the ChAT^+^Islet1^-^ are indeed interneurons and not motoneurons which had lost the GFP expression in the *Islet1* line, we injected a retrograde tracer into zebrafish spinal cord ventral roots (see Material and Methods; Figure 1H) and combined it with ChAT immunoreactivity (Figure 1I). Given that all zebrafish motoneuron axons exiting from the same ventral root correspond to the spinal hemisegment, this approach enabled us to reveal all the spinal motoneurons (axial and fin; Figure 1H). We observed that 37.37 ± 3.75% (Figure 1J) of the cholinergic neurons are not motoneurons, similar to findings in the *Islet1* zebrafish transgenic line.

Finally, to assess whether spinal cholinergic interneurons innervate targets located in rostral or caudal spinal cord segments, we identified cholinergic interneurons (into segment 15) as ascending or descending, after injection of a retrograde tracer (dextran) into segments 12 or 18, respectively, in an *Islet1:GFP* line (n=14 zebrafish; Figure 1J,K). 24h after the tracer injection we processed the tissue for ChAT immunostaining. We observed that 7.56% of ChAT^+^Islet1^-^ neurons were labelled as descending neurons (Supplementary file 1J, L, M), and 13.22% of ChAT^+^Islet1^-^ neurons were determined as ascending neurons (Supplementary file 1K-M), indicating that 20.78% of the cholinergic interneurons project to other spinal cord segments. The significant difference between the soma size of the ascending and descending cholinergic interneurons suggest that they form two non-overlapping neuronal subpopulations (Supplementary file 1N). Overall, our data suggest that zebrafish spinal motoneurons receive unevenly distributed cholinergic input from a local spinal source, represented by cholinergic interneurons.

### V0c interneurons are present in zebrafish spinal cord

Most (~75%) ChAT interneurons (ChAT^+^Islet1^-^) were found in the dorsal part of the motor column close to the central canal (Figure 1G), where the mammalian V0c interneurons are also positioned (Zagoraiou et al., 2009). Previously, it was assumed that V0c interneurons do not exist in zebrafish (Satou et al., 2012). To test if the observed zebrafish spinal cholinergic interneurons correspond to the mammalian V0c interneurons, we combined *in situ* hybridization for *Pitx2,* a paired-like homeodomain transcription factor that marks V0c interneurons in mammals (Figure 2A; Zagoraiou et al. 2009), with ChAT immunolabeling (Figure 2B). Our analysis revealed that 28.36 ± 1.83 % of the ChAT neurons were also *Pitx2* positive (*Pitx2^+^*; Figure 2C). Although the *Pitx2^+^* neurons were distributed throughout the motor column, ~80% were located around the central canal (Figure 2D), suggesting that an equivalent of mammalian V0c interneurons do exist in zebrafish. Moreover, we performed whole cell patch clamp recordings from several unidentified neurons located around the central canal. We found that 3 neurons out of 20 were cholinergic and not motoneurons, and they displayed intrinsic electrical properties (i.e. prominent after-hyperpolarization, rheobase, lack of neuronal sag) comparable to the mammalian V0c interneurons (Zagoraiou et al. 2009; Figure 2E). Although these interneurons do not receive rhythmic synaptic inputs during locomotion, they receive strong synaptic potentials at speeds < 6 Hz (paired t test, t = 5.133 df = 40, p < 0.0001, n = 3 neurons; Figure 2F).

**Figure 2.**
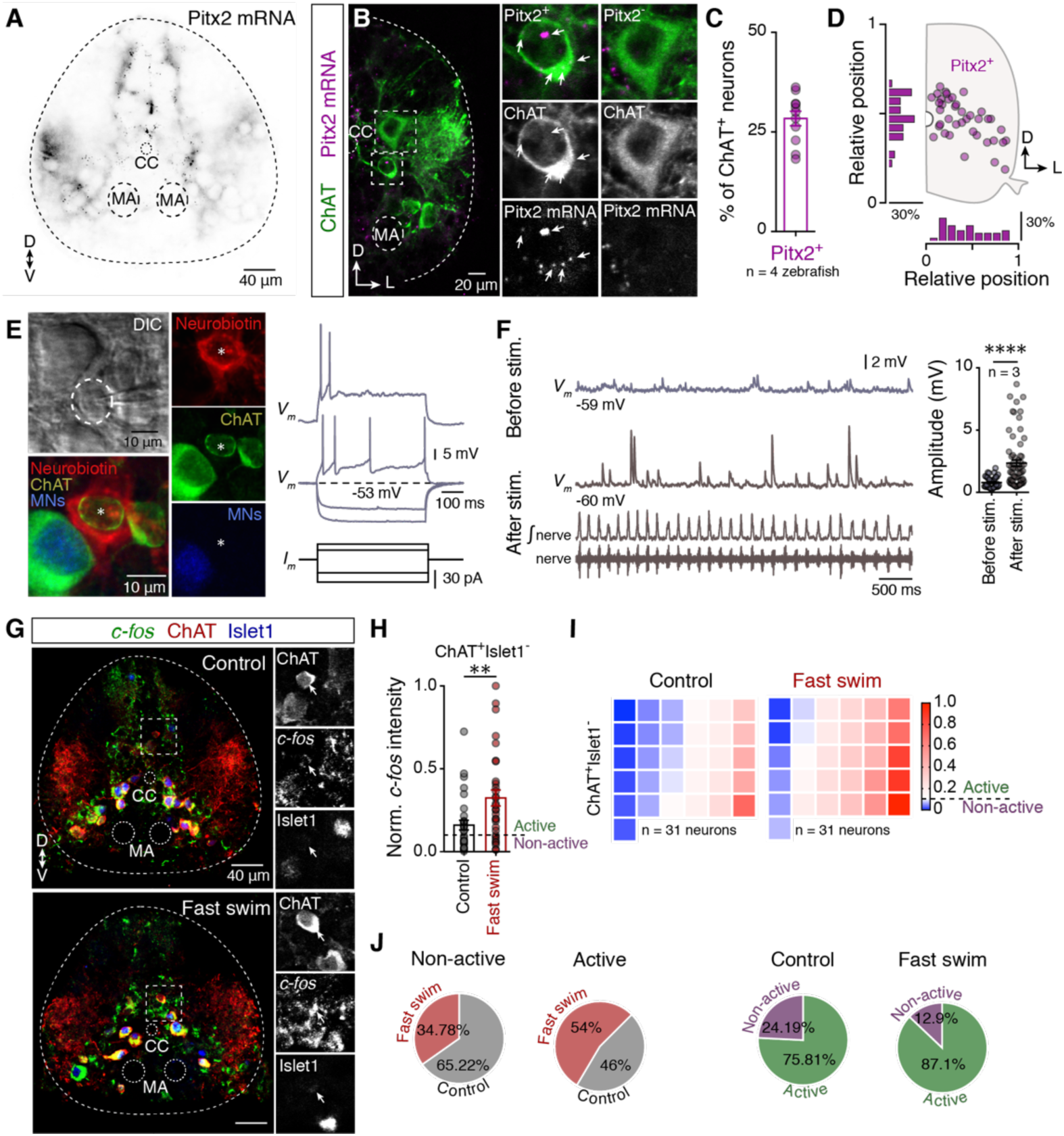
Cholinergic interneurons are homologues to mammalian V0c interneurons and they are recruited at high speed locomotion. (**A**) Localization of *Pitx2 mRNA* in adult zebrafish spinal cord. (**B**) Cholinergic *Pitx2^+^* and *Pitx2^-^*neurons found in adult zebrafish spinal cord. Arrows indicate the location of mRNA in the cholinergic neuron. (**C**) Quantification (%) of cholinergic neurons that are *Pitx2^+^* (n = 4 zebrafish). (**D**) Distribution pattern of cholinergic *Pitx2^+^* interneurons within the adult zebrafish spinal cord. (**E**) Whole cell patch clamp recording from identified neuron that filled with neurobiotin. Post-hoc evaluation of the patched neuron that is cholinergic (ChAT^+^) and not motoneuron (MN^-^). Firing behavior of a representative cholinergic interneuron upon steps of somatic current injections. The asterisk denotes the neurobiotin filled patched neuron. (**F**) Increasing number and amplitude of post-synaptic responses during fictive locomotion, as indicated from the recording of motor nerve activity (lower trace and rectified trace) after electrical stimulation of descending axons to initiate locomotion (n = 3 neurons). (**G**) Microphotographs showing that *c-fos* immunoreactivity increases after prolonged (2h) forced fast swimming (80% of the Ucrit) in the cholinergic interneurons (ChAT^+^Islet1^-^). (**H**) Quantification of normalized *c-fos* intensity in cholinergic interneurons (ChAT^+^Islet1^-^). The threshold of activity was considered to be 0.11. (**I**) Heat map displaying the normalized *c-fos* intensity of each individual cholinergic interneuron (n_control_ = 31 neurons, n_fast swim_ = 31 neurons). (**J**) Proportions of active and non-active cholinergic interneurons (ChAT^+^Islet1^-^) between control and after forced fast swimming based on the normalized *c-fos* intensity. CC, central canal; D, dorsal; L, lateral; MA, Mauthner axon; P, posterior; V, ventral. Data are presented as mean ± SEM; **p < 0.01; ****p < 0.0001.

It was apparent that the cholinergic interneurons are not active during regular swimming (< 6Hz), so we next assessed when these cholinergic interneurons operate, contributing to locomotor network functionality. To address this issue, we subjected adult zebrafish to forced fast swimming test (80% of the critical speed (Ucrit), see Material and Methods) for 2 h, followed by functional anatomical analysis of *c-fos* expression as an index of neuronal activity. Analysis of *c-fos* intensity in all ChAT^+^Islet1^-^ neurons revealed that more neurons are active (54%) during fast swimming, compared to the control, as indicated by a significant increase in *c-fos* expression (unpaired t test, t = 2.89 df = 60, p = 0.0054; Figure 2G-J). Our observations are in agreement with previous analysis performed in the cat spinal cord, which showed that cholinergic neurons close to the central canal were functionally active during prolonged fictive locomotion (Huang et al., 2000). Collectively, our results demonstrate that the vast majority of the zebrafish cholinergic interneurons (80%) are analogues to mammalian V0c interneurons and their activity is related to prolonged fast locomotion.

**Muscarinic receptors differentially alter motoneuron excitability**

In mammals, acetylcholine increases the excitability of spinal motoneurons through the activation of muscarinic receptors (Miles et al., 2007). To investigate whether zebrafish motoneuron excitability is responsive to activation of mAChRs we obtained whole cell current clamp recordings from different classes (slow, intermediate and fast) of axial secondary motoneurons. Since the fast and intermediate motoneurons discharge single action potentials (APs), whereas slow motoneurons fire in bursts of APs (Gabriel et al., 2011; Ampatzis et al., 2013), the excitability of motoneurons was assessed by examining their response (number of single APs or number of bursts) to steps of supra-threshold depolarizing current pulses (500 ms, increments of 10% from rheobase), before and after the application of muscarine, a non-selective mAChR agonist. In the presence of muscarine (15 μM), the firing rate of both slow (one-way ANOVA repeated measures, F_1.319, 5.277_ = 16, p = 0.021, n = 5 out of 5) and intermediate (one-way ANOVA repeated measures, F_1.269, 6.344_= 24.36, p = 0.0079, n = 6 out of 6 neurons) motoneurons substantially increased (Figure 3A-C, F). This was accompanied by an increase of the input resistance (slow MNs: one-way ANOVA repeated measures, F_1.059, 4.237_ = 13.34, p = 0.0227; intermediate MNs: one-way ANOVA repeated measures, F_1.493, 7.466_ = 12.64, p = 0.0258; Figure 3E) and changes in other intrinsic biophysical properties (Supplementary file 2), without significantly altering the resting membrane potential (Figure 3D). In contrast, the fast motoneurons showed a decrease in excitability, resulting in fewer action potentials (one-way ANOVA repeated measures, F_1.986, 15.89_ = 38.05, p < 0.0001, n = 9 out of 9 neurons; Figure 3A,B,F) associated with a significant decrease of the input resistance (one-way ANOVA repeated measures, F_1.12, 8.958_ = 4.462, p = 0.0141, Figure 3E). Moreover, alterations of the electrical properties of fast motoneurons were similar to those observed in the intermediate motoneurons (Supplementary file 2). The observed changes in intrinsic biophysical properties of the motoneurons are consistent with previously reported alterations of mammalian motoneurons in response to muscarine (Ireland et al., 2012).

**Figure 3.**
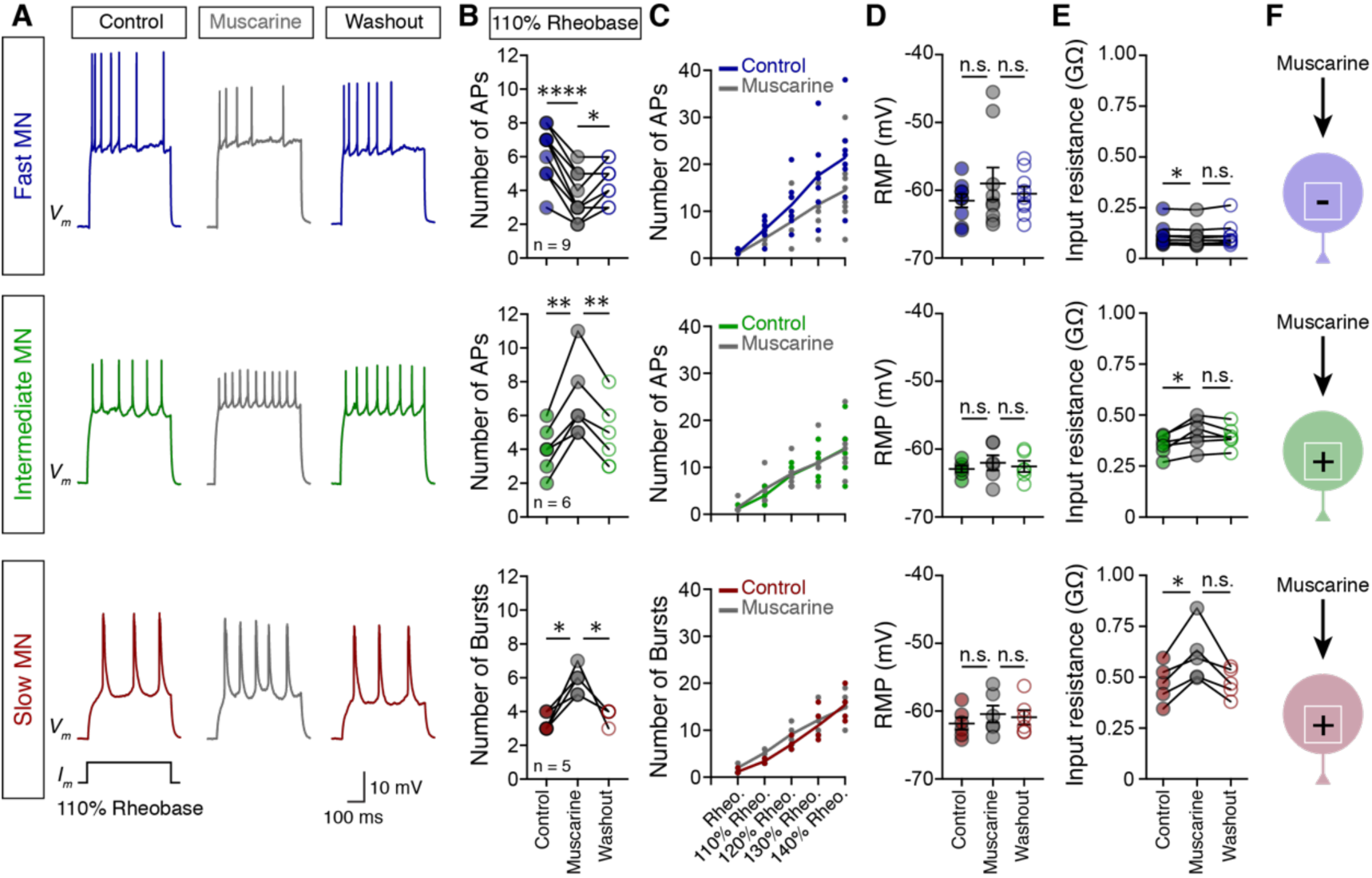
Muscarinic receptors differentially alter motoneuron excitability. (**A**) Representative traces of motoneuron (MN) firing in responses to 110% of rheobase somatic current injection, before (colored traces) after bath application of muscarine (15 μM; gray traces) and followed by washout (colored traces). (**B**) In the presence of muscarine the fast motoneurons significantly reduce the number of action potentials (APs) in response to 110% of rheobase current injection, while both the intermediate and the slow motoneurons increase their firing rate. During washout, the motoneurons partially recover (n_fast_ = 9 neurons, n_intermediate_ = 6 neurons, n_slow_ = 5 neurons). (**C**) Relation of the number of action potentials (APs) or bursts of APs to current (increments of 10% increase from rheobase) in absence (colored line) and presence of muscarine (gray line). (**D**) During the application of muscarine (gray circles) and after washout (open colored circles) there is no significant change in the resting membrane potential of the motoneurons. (**E**) Bath application of muscarine alters the input resistance of all motoneuron types. The input resistance was decreased in the fast motoneurons and increased in the intermediate and slow motoneurons. (**F**) Overview of the change in excitability of different types of MNs in response to muscarine. The sign represents the alteration of the MN excitability (+ = increase, - = decrease). Data are presented as mean ± SEM; *p < 0.05; **p < 0.01; ****p < 0.0001; n.s., non-significant.

### Motoneurons possess different muscarinic receptor subtypes

We speculated that the differential motoneuron excitability we observed might arise from the presence of different mAChR subtypes in the motoneurons. To test if m2-mAChRs alone mediate the changes in motoneuron excitability, we applied a mixture of muscarine (15 μM) and methoctramine (10 μM), a selective m2-mAChR antagonist (Hulme et al., 1990). We recorded an increase in excitability of fast (paired t test, t = 3.641, p = 0.0219, n = 5 out of 5 neurons; Figure 4A-C), intermediate (paired t test, t = 4.323, p = 0.0228, n = 4 out of 4 neurons) and slow (paired t test, t = 4.00, p = 0.0161, n = 3 out of 3 neurons) motoneurons (Figure 4A-C). To assess whether m2-mAChRs are present in all classes of motoneurons, we applied oxotremorine-M (Oxo-M; 20 μμ), an m2-mAChR preferential agonist (Murakami et al., 1996). In response, the fast (paired t test, t = 10.00, p = 0.0005, n = 5 out of 5 neurons) and intermediate (paired t test, t = 6.668, p = 0.0026, n = 5 out of 5 neurons) motoneurons exhibited reduced firing (Figure 4D-F), whereas, slow motoneuron firing was unaffected (paired t test, t = 1.00, p = 0.391, n = 4 out of 4 neurons; Figure 4D-F), suggesting that they do not express the m2-mAChR subtype. Moreover, we confirmed the presence of m2-mAChRs in motoneurons anatomically. Immunoreactivity for the m2-mAChRs was strong in fast motoneurons and modest in intermediate motoneurons (One-way ANOVA, F_2,35_ = 51.32, p < 0.0001, n = 38 neurons; Figure 4G, H). In contrast, slow motoneurons were weakly labeled for the m2-mAChR, supporting previous reports (Welton et al., 1999; Figure 4G, H). These results indicate that multiple mAChR subtypes are expressed in a motoneuron type-dependent manner, which functions to differentially mediate the actions of acetylcholine.

**Figure 4.**
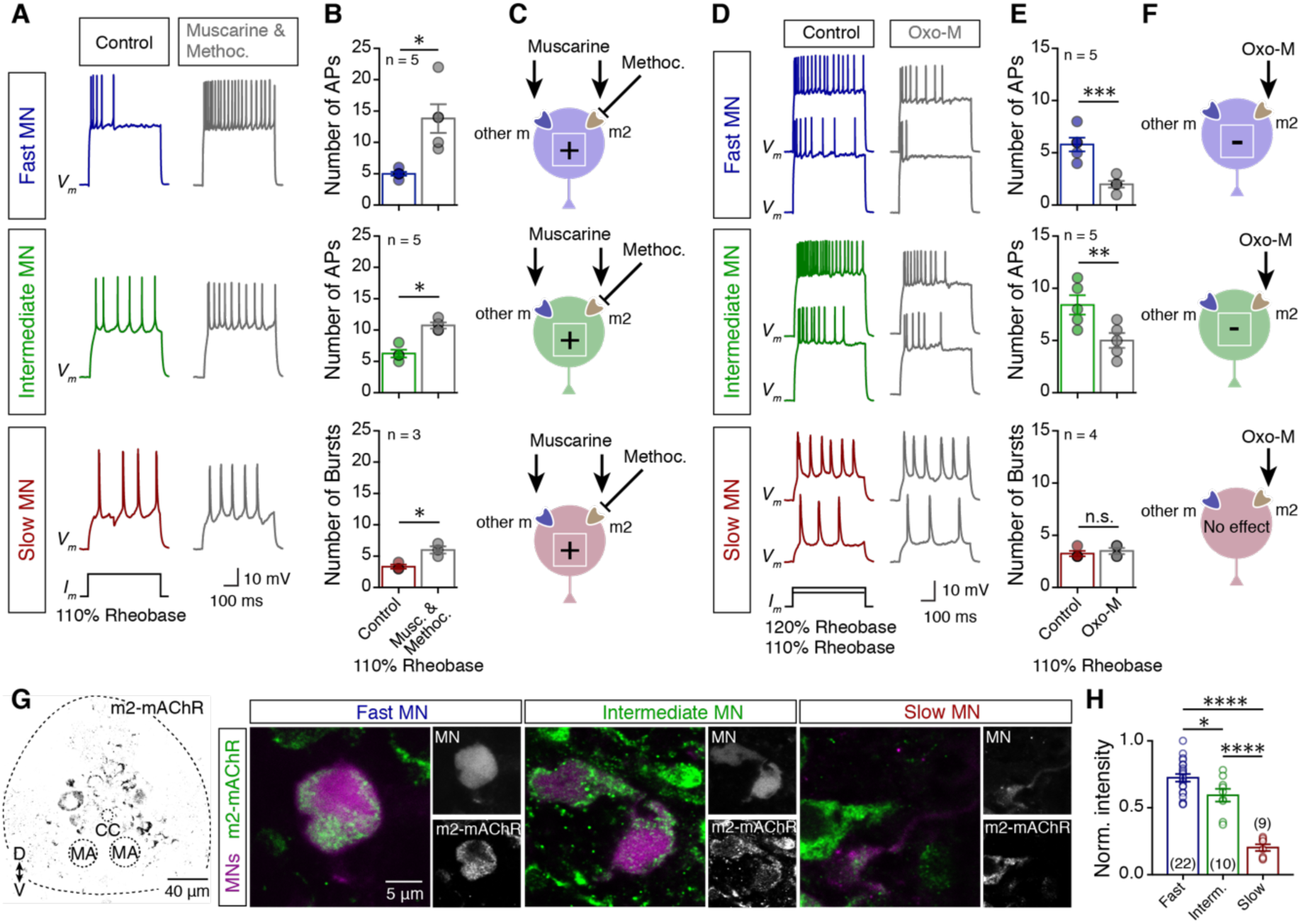
Motoneurons possess different muscarinic receptor subtypes. (**A**) Traces of motoneuron (MN) firing in response to 110% of rheobase current injection, before (colored traces) and after bath application of a mixture of muscarine (15μM) and methoctramine (10μM; gray traces). (**B-C**) Co-application of muscarine and methoctramine produce hyperexcitability in all MN pools (n_fast_ = 5 neurons, n_intermediate_ = 5 neurons, n_slow_ = 3 neurons), derived from the antagonistic effect on m2-mAChRs. (**D**) Representative traces of motoneuron (MN) firing in response to 110% and 120% of rheobase current injections, before (colored traces) and after bath application of oxotremorine-M (Oxo-M; 20 μμ; gray traces) which preferentially activates the m2-mAChRs. (**E-F**) In the presence of oxotremorine-M, excitability decreases in the fast and intermediate motoneurons, whereas the slow motoneurons do not change their firing rate in response to 110% of rheobase current injection (n_fast_ = 5 neurons, n_intermediate_ = 5 neurons, n_slow_ = 4 neurons). (**G**) Distribution pattern of m2-mAChRs in relation to different motoneuron types. A combination of retrograde labeling of different motoneuron pools with immunostaining for m2-mAChRs reveals that the fast motoneurons exhibit a vast number of m2 receptors compared to other motoneuron types (intermediate and slow). Slow motoneurons were found to be weakly labeled. The sign represents the excitability alteration of the MN (+ = increase, - = decrease). Data are presented as mean ± SEM; *p < 0.05; **p < 0.01; ***p < 0.001; ****p < 0.0001; n.s., non-significant.

### Activation of motoneuron muscarinic receptors controls fast locomotion

Given that motoneuron excitability can retrogradely influence the upstream locomotor network function (presynaptic release, generation of action potentials, swimming duration and frequency) through gap junctions (Song et al., 2016), we tested the functional significance of motoneuron mAChR activation during locomotion. For this, we used the adult zebrafish *ex vivo* preparation (Ampatzis et al., 2013; Ampatzis et al., 2014; Gabriel et al., 2011; Song et al., 2016). Bath application of muscarine facilitated fictive locomotor activity and increased the highest reached swimming frequency by 26.8 ± 5.8 % (paired t test, t = 6.675, p < 0.0001, n = 12; Figure 5A-B). It should be noted that the amplitude of the swimming membrane potential oscillations during locomotion is related to swimming frequency (Gabriel et al., 2011). In the presence of muscarine, the correlation between oscillation amplitude and frequency was decreased in the fast motoneurons (Figure 5C) and enhanced in the intermediate motoneurons that were recruited above 7 Hz (Figure 5D). Furthermore, we investigated the number of APs in the slow and intermediate motoneurons during ongoing fictive locomotion at 5 Hz (Figure 5E). At this swimming speed, we found that only the intermediate motoneurons discharge more action potentials in the presence of muscarine, compared to controls (paired t test, t = 3.43, p = 0.0018; Figure 5E). Overall, our findings demonstrate that activation of spinal muscarinic receptors facilitates the recruitment of the intermediate motoneuron-interneuron module while hindering the recruitment of the fast locomotor module, as seen through the motoneuron activation.

**Figure 5.**
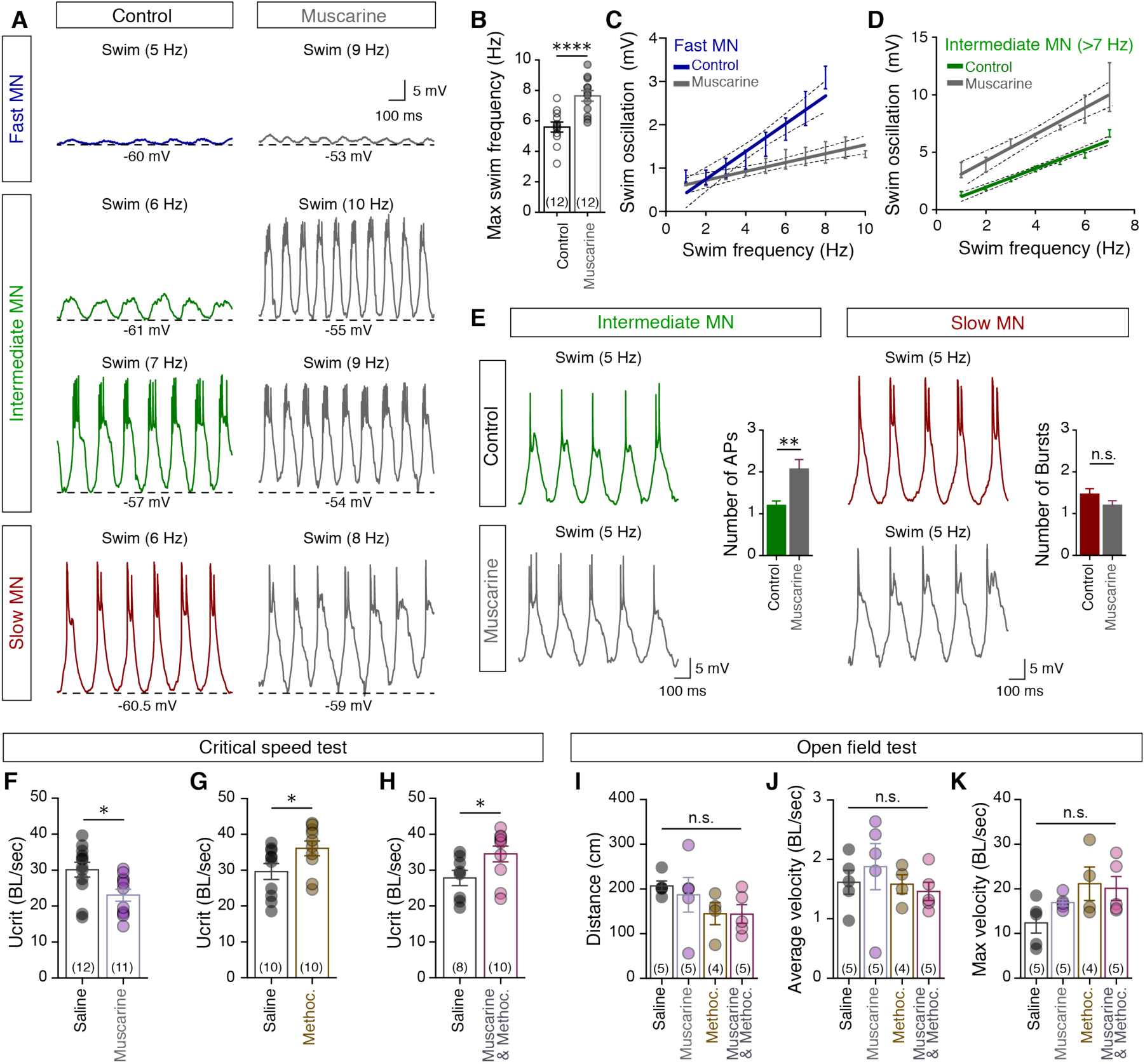
Activation of motoneuron muscarinic receptors controls fast locomotion. (**A-B**) During fictive locomotion, motoneurons display membrane potential oscillations. The highest swimming frequency increases in the presence of muscarine (15 μM; gray traces) during fictive locomotion (*ex vivo* experiments). Dashed line indicates the membrane potential of the lower part of the oscillations. (**C**) Averaged slopes of the membrane potential oscillations in relation to different swimming frequencies of fast motoneurons in the absence (blue line), or presence (gray line) of muscarine. The data are presented as interpolation curve (middle line ± SEM) and 95% confidence band (outer dashed lines).(**D**) Slopes showing the increasing amplitude of the membrane potential oscillations of the intermediate motoneurons recruited above 7 Hz, in relation to the swimming frequency, before and after the application of muscarine. After muscarine (gray line) the amplitude of the membrane oscillations was significant increased (F = 856.6 p < 0.0001), in comparison to the control (green line). The data are presented as interpolation curve (middle line ± SEM) and 95% confidence band (outer dashed lines). (**E**) During 5 Hz fictive locomotion, in the absence (colored trace) or presence (gray trace) of muscarine, the number of discharges of intermediate motoneurons increased, while slow motoneurons did not display any changes in the number of bursts of action potentials. (**F-H**) Intraperitoneal administration of muscarine reduces the U_crit_ (BL = 1.84 ± 0.11 cm; n = 23 zebrafish), while methoctramine increased the maximum swimming speed (BL = 1.68 ± 0.1 cm; n = 20 zebrafish). Co-administration of muscarine and methoctramine increased the maximum obtained swimming speed (BL = 1.71 ± 0.12 cm; n = 18 zebrafish). (**I-K**) *In vivo* monitoring of adult zebrafish locomotor behavior. Muscarine and/or methoctramine administration was not found to affect the distance traveled, the average velocity and the maximum velocity of the studied animals (n = 19 zebrafish). The data were normalized to body lengths (BL) / sec. Data are presented as mean ± SEM; *p < 0.05; **p < 0.01; ****p < 0.0001; n.s., non-significant.

Finally, we sought to better understand the *in vivo* behavioral functions of the activation of mAChRs. Therefore, we subjected zebrafish to a critical speed test (Supplementary file 3; see Supplemental Experimental Procedures). Critical speed (U_crit_) is a measure of the highest sustainable swimming speed that a fish can reach (Brett, 1964). Intraperitoneal administration of muscarine (50 μM, 525 ng/g BW; Supplementary file 3A) significantly reduced the highest sustainable swimming speed (unpaired t test, t = 2.65, p = 0.014, n = 23 zebrafish; Figure 5F).

In contrast, methoctramine treated animals (40 μM, 2.2mg/g BW; Supplementary file 3) increased the uppermost locomotor speed (unpaired t test, t = 2.12, p = 0.047, n = 20 zebrafish; Figure 5G). Similarly, exogenous activation of all muscarinic receptors except the m2-mAChRs, achieved by co-administration of muscarine (50 μM) and methoctramine (40 μM), increased the maximum locomotor speed obtained (unpaired t test, t = 2.156, p = 0.046, n = 18 zebrafish; Figure 5H).

To test the effect of our pharmacological treatments on locomotion under normal conditions, animals were subjected to an open field test (Supplementary file 3, see Material and Methods). We observed that similar treatment did not affect the regular locomotor behavior (distance traveled, average velocity and maximum velocity; Figure 5I-K), suggesting that the differences in critical speed observed here are due to the effect on the spinal locomotor circuit. These data show that stimulation of m2-mAChRs causes a reduction in the maximum swimming speed to 76.5 ± 6.7 % of the of the U_crit_, corresponding to the zebrafish optimum speed (Palstra et al., 2010), where swimming relies only on the activity of the slow and intermediate neuro-muscular system, and not on the fast. Our analysis cannot rule out the possibility of an additional direct effect of mAChR activation in the premotor neurons, however, as any alteration in motoneuron excitability will retrogradely affect the excitability and pre-synaptic release of glutamate from V2a interneurons (Song et al., 2016), which will influence premotor network functionality. Taken together, our results suggest that cholinergic modulation acts both on motoneurons and the premotor network to modify the speed of locomotion mainly through the engagement of the fast motoneuron-interneuron module.

## Discussion

Our findings suggest that motoneurons receive cholinergic input exclusively from spinal interneurons, with the vast majority of them being analogous to mammalian V0c interneurons. Acetylcholine shifts motoneuron excitability through the parallel activation of different mAChRs subtypes (Figure 6A). Moreover, activation of this pathway selectively alters the operational range of different motoneuron modules during locomotion (Figure 6B). Overall, this work provides novel insights into how intraspinal acetylcholine release can modify the functionality of the spinal circuitry, which requires synaptic specificity and temporal precision to generate locomotion at different speeds (Ampatzis et al., 2013; Ampatzis et al., 2014).

**Figure 6.**
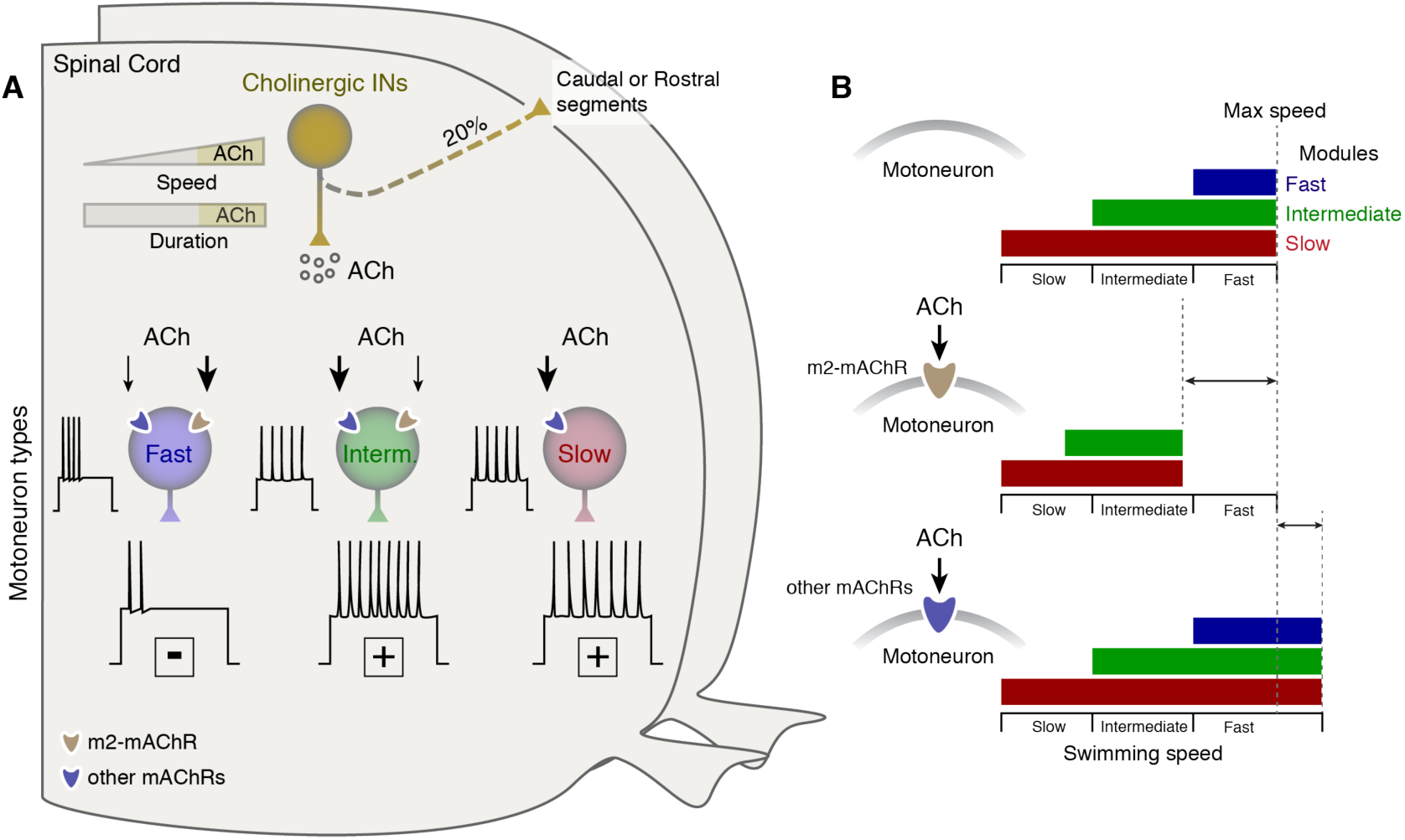
Differential organization of cholinergic input to MNs permits changes of the operational range of locomotor circuit. (**A**) Acetylcholine release (after prolonged fast swimming) from spinal cholinergic interneurons affects the excitability of different types of motoneurons in a mAChR dependent manner. Our results clearly demonstrate that acetylcholine can only increase slow motoneuron excitability, whereas the intermediate and fast motoneurons can alter their excitability through the activation of the different mAChRs that they possess. (**B**) This configuration allows the animals to adjust the recruitment of the locomotor microcircuit modules and alter the operational range of these networks.

One of our major findings is establishing the existence of cholinergic interneurons in the zebrafish spinal cord and identifying them as the exclusive source of the cholinergic spinal input. Although earlier studies showed the existence of cholinergic interneurons in several vertebrate species (Thiriet et al., 1992; Anadón et al., 2000; González et al., 2002; Miles et al., 2007; Quinlan and Buchanan, 2008; Zagoraiou et al., 2009), and moreover, the zebrafish cholinergic system is well characterized (Clemente et al., 2004; Mueller et al., 2004), the presence of zebrafish spinal cord cholinergic interneurons has not been previously reported. While a number of Chat^+^Islet1^-^ neurons were identified earlier (Stil and Drapeau, 2016), it has been proposed that these small cholinergic neurons correspond to motoneurons, rather than the ones that innervate the axial muscles (Muller et al., 2004). Previous attempts to localize *Pitx2* expression with *Dbx1* in order to reveal the existence of V0c interneurons in zebrafish spinal cord were ineffective (Satou et al., 2012). In the current study we used a different approach, by combining *Pitx2* expression with ChAT immunostaining, to achieve this goal. We revealed that most cholinergic interneurons share comparable anatomical distribution, molecular identity and electrical properties with mammalian V0c interneurons. Our analysis cannot rule out the possibility that more cholinergic interneuron subtypes are present in zebrafish spinal cord. However, most of zebrafish cholinergic interneurons (~80%) express *Pitx2*, thus we considered them to be similar to mammalian V0c interneurons. Since V0c interneurons have been shown to be the exclusive source of cholinergic c-bouton input to mammalian motoneurons (Zagoraiou et al. 2009), our data directly implicate the activity of these cholinergic interneurons in the cholinergic modulation of motoneurons in the zebrafish.

The overall activation of the muscarinic receptors has been shown to increase motoneuron excitability (Alaburda et al., 2002; Chevallier et al., 2006; Hornby et al., 2002; Ireland et al., 2012; Miles et al., 2007) via activation of the m2-mAChR subtype (Miles et al., 2007). On the other hand, numerous studies have shown that activity of the m2-mAChRs is associated with inhibitory actions in the nervous system (Brown, 2010; Felder, 1995; Hosey, 1992; Kurihara et al., 1993). Our findings challenge previous conclusions that activation of motoneuron m2-mAChR is primarily required to increase motoneuron excitability. By pharmacologically manipulating the m2-mAChRs in different motoneuron pools, we found that m2-mAChR activation reduces motoneuron excitability and other receptor subtypes can account for the observed rise in motoneuron firing. In line with this idea, various subtypes of mAChRs have been characterized in the spinal cord neurons (Brown, 2010; Höglund and Baghdoyan, 1997; Zhang et al., 2005; Jordan et al., 2014) and it is also well documented that many nerve cells contain more than one subtype of muscarinic receptor (Finkel et al., 2014; Hassall et al. 1993; Jordan et al., 2014). The exact muscarinic receptor subtype responsible for the elevated excitability of adult zebrafish motoneurons remains to be identified. However, it has been suggested that the excitatory effects of muscarinic agonists on neonatal rat motoneurons are mediated through the m3-mAChRs (Kurihara et al., 1993). In support of our observations, Jordan et al. (2014) recently demonstrated that m2-mAChRs and m3-mAChRs are both involved in the cholinergic modulation of locomotion in mammals. They showed that application of the m2-mAChR antagonist methoctramine, increased the locomotor rhythm, whereas application of m3-mAChR antagonist reduced, and finally blocked, the locomotor activity (Jordan et al., 2014). Our results show that different muscarinic receptor subtypes in the membrane of motoneurons can play a unique role in the regulation of neural activity involved in the control of locomotion.

Given the emerging evidence regarding the importance and functional repertoire of spinal cholinergic input in the control of motor behaviors, it is not surprising that cholinergic synapses have been implicated in spinal cord injury (Ichiyama et al., 2011; Skup et al., 2012; Jordan et al., 2014) and motor disease (Herron and Miles, 2012; Pullen and Athanasiou, 2009; Saxena et al., 2013). However, no causal link has yet been found between alterations in the number and size of cholinergic synapses, acetylcholine release and muscarinic receptor activation. Therefore, uncovering the mechanisms by which acetylcholine modifies the activity of spinal neurons is of significant biological and medical interest. We find that methoctramine treated animals are able to operate at higher swimming speeds and, in addition, activation of m2-mAChRs reduces the excitability of the motoneurons, and potentially of other spinal cord neurons, and their ability to produce the appropriate muscle force. In support of our findings, Jordan et al., (2014) revealed that application of methoctramine increased the locomotor frequency in mammals. Moreover, application of cholinergic receptor antagonists following spinal cord injury was found to advance locomotor activity, suggesting that adaptive alterations of the spinal cholinergic system obstruct the generation and execution of locomotion (Jordan et al., 2014). In a mouse model of amyotrophic lateral sclerosis (ALS), methoctramine treatment counteracts the loss of muscle force during the pre-symptomatic stages of the disease (Saxena et al., 2013), and is likely to be mediated through the blocking of m2-mAChRs, increasing the excitability of spinal neurons participating in the generation of movement. In conclusion, our results provide a novel contribution to existing knowledge, and further understanding of the causal relationship between activation of muscarinic receptors and locomotion, highlighting their potential as targets for innovative therapeutic strategies.

A general question about the cholinergic control of locomotion is made addressable in our work: how does the muscarinic acetylcholine receptor configuration gate the execution of locomotor behavior at different speeds? It has been proposed that the cholinergic modulation of motoneurons is activity dependent (Zagoraiou et al., 2009). In favor of this idea, our work revealed that pharmacological manipulations of spinal circuit muscarinic receptors *ex vivo* and *in vivo* only affect the locomotor performance of zebrafish at higher swimming speeds, which are primarily associated with the engagement and function of the fast motoneuron-interneuron module. Thus, the mechanisms we identify here suggest an additional level of control during high-energy-demand locomotion, such as fast swimming.

### Materials and Methods

#### Animals

All animals were raised and kept in a core facility at the Karolinska Institute according to established procedures. Adult zebrafish (*Danio rerio*; 8-10 weeks old; length, 17-19 mm; weight, 0.025-0.045 g) wild type (AB/Tübingen) and Tg(*Islet1:GFP*) lines where used in this study. All experimental protocols were approved by the local Animal Research Ethical Committee, Stockholm and were performed in accordance with EU guidelines.

#### Motoneuron and descending/ascending neuron labeling

Zebrafish (n = 30) of either sex were anaesthetized in 0.03% tricaine methane sulfonate (MS-222, Sigma-Aldrich). Retrograde labeling of axial motoneurons was performed using dye injections with tetramethylrhodamine-dextran (3000 MW; ThermoFisher, D3307) in specific muscle fiber types (slow, intermediate or fast). In addition, retrograde labeling of all motoneurons was performed using similar procedure to spinal cord ventral roots. To label the neurons descending from the brain to the spinal cord the tracer was injected using dye-soaked pins in the spinal cord at approximately the level of the 6-8th vertebra. Finally, for the investigation of descending and ascending cholinergic interneurons the tracer was injected in the spinal segment 18 (n = 7 zebrafish) or 12 (n = 7 zebrafish) respectively. Afterwards all the animals were kept for at least 24h to allow the retrograde transport of the tracer. We evaluated the number of ChAT^+^Istlet1^-^ ineterneurons that were positive to the tracer on segment 15 of the adult zebrafish spinal cord.

#### Immunohistochemistry

All animals were deeply anesthetized with 0.1% MS-222. We then dissected the spinal cords and/or the brains and fixed them in 4% paraformaldehyde (PFA) in phosphate buffer saline (PBS) (0.01M; pH = 7.4) at 4°C for 2-14h. We performed immunolabelings in both whole mount and cryosections. For cryosections, the tissues were removed carefully and cryoprotected overnight in 30% (w/v) sucrose in PBS at 4°C, embedded in OCT Cryomount (Histolab), rapidly frozen in dry-ice-cooled isopentane (2-methylbutane; Sigma) at approximately –35°C, and stored at –80°C until use. Transverse coronal plane cryosections (thickness 25 μm) of the tissue were collected and processed for immunohistochemistry. The tissue was washed three times for 5 min in PBS. Nonspecific protein binding sites were blocked with 4% normal donkey serum with 1% bovine serum albumin (BSA; Sigma) and 0.5% Triton X-100 (Sigma) in PBS for 30 min at room temperature (RT). Primary antibodies were diluted in 1% of blocking solution and applied for 24-90 hours at 4 °C. For primary antibodies, we used goat anti-ChAT (1:200; Millipore, AB144P, RRID: AB_2079751), rabbit anti-GFP (1:700; ThermoFisher Scientific, A-11122, RRID: AB_221569), guinea pig anti-VAChT (1:1000; Millipore, AB1588, RRID: AB_11214110), rabbit anti-cfos (1:200; Sigma, F7799) and rabbit anti-m2 mAChRs (1:700-1:2000; Alomone, AMR-002, RRID: AB_2039995). After thorough buffer rinses the tissues were then incubated with the appropriate secondary antibodies diluted 1:500 in 0.5% Triton X-100 (Sigma) in PBS overnight at 4 °C. We used Alexa Fluor–conjugated secondary antibodies anti-goat 488 (ThermoFisher Scientific, A11055, RRID: AB_142672), anti-goat 568 (ThermoFisher Scientific, A11057, RRID: AB_142581), anti-rabbit 488 (ThermoFisher Scientific, A21206, RRID: AB_141708) and anti-guinea pig 488 (ThermoFisher Scientific, A11073, RRID: AB_142018). Finally, the tissues were thoroughly rinsed in PBS and cover-slipped with fluorescent hard medium (VectorLabs; H-1400).

#### In situ hybridization

Adult zebrafish wild type (AB/Tübingen) spinal cords were fixed, cryoprotected and cryosectioned (thickness of 12 μm) as described before (see *Immunohistochemistry* section). Coronal transverse section then processed for fluorescent *in situ* hybridization using the RNAscope Technology (Advance Cell Diagnostics), according to manufacturer’s instructions. Target probe for zebrafish specific *Pitx2* gene (Advance Cell Diagnostics, 432161-C3) was used. After the detection of *Pitx2* mRNA, tissues were processed for ChAT immunodetection as described before (*Immunohistochemistry section*). To investigate how many of cholinergic neurons are *Pitx2*, 3 sections per animal (n = 4 zebrafish) corresponding to 12-14th spinal cord segments were analyzed.

#### *c-fos* functional anatomy

Adult zebrafish (*Islet1:GFP*; n = 8) were used for c-fos experiments. Animals were subjected to forced swim test. We used two different experimental protocols to investigate the activity of cholinergic interneurons. Animals were subjected to prolonged swimming (2 h) at 80% of the critical speed (U_crit_). Control animals were kept under regular tank swimming conditions. Immediately after the test all animals were fixed and processed for triple immunolabeling, against *c-fos*, the ChAT and the GFP (see Supplemental Experimental Procedures, section Immunohistochemistry) in 25 μm thick cryosections. For the *c-fos* experiments, analysis was performed in neurons (n=62) from sections corresponding to segments 14-17. The intensity of the *c-fos* activity was evaluated in the obtained confocal pictures using ImageJ.

#### Microscopy and image analysis

Imaging was carried out in a laser scanning confocal microscope (LSM 510 Meta, Zeiss). Cholinergic inputs on different motoneuron types were counted on single plan confocal images. Counting was performed in non-overlapping fields of spinal cord sections, in spinal cord segments 14-17. We defined the putative cholinergic inputs as number of large (>0.5 μm of diameter) VAChT-positive putatively cholinergic synapses, apposing the somata of motoneurons. The evaluation included 10 slow, 27 intermediate and 30 fast positive motoneurons collected from 4 adult zebrafish spinal cords. For the quantification of m2-mAChRs, all images were captured at identical exposure times in order to ensure the same illumination level. The intensity of m2-mAChRs immunoreactivity was evaluated in the obtained confocal pictures using the ImageJ image analysis software. The relative position of the somata of the neurons within spinal cord, was calculated in whole mount preparations, using the lateral, dorsal, and ventral edges of spinal cord as well as the central canal as landmarks. Analysis of all spinal cord neurons was performed between segments 14-17. The relative position was calculated using ImageJ. Examination of the descending neurons was performed from a series of coronal brain section, throughout the brain, without discarding any section from the analysis. The nomenclature used for the brain areas of descending neurons was based on the topological zebrafish brain atlas (Wullimann et al., 1996). All figures and graphs were prepared with Adobe Photoshop and Adobe Illustrator (Adobe Systems Inc., San Jose, CA). All double-labeled images were converted to magenta-green immunofluoresence to make this work more accessible to the red-green color-blind readers.

#### *Ex vivo* preparation and electrophysiology

The dissection procedure has been described previously (Gabriel et al., 2011; Ampatzis et al., 2013; Ampatzis et al., 2014; Song et al., 2016). The preparations (n = 47) were then transferred to a recording chamber, placed lateral side up, and fixed with Vaseline. The chamber was continuously perfused with extracellular solution contained: 134 mM NaCl, 2.9 mM KCl, 2.1 mM CaCl_2_, 1.2 mM MgCl_2_, 10 mM HEPES, and 10 mM glucose, pH 7.8, adjusted with NaOH, and an osmolarity of 290 mOsm. All experiments were performed at an ambient temperature of 20–22°C. For whole-cell intracellular recordings, electrodes (resistance, 9–13 MO) were pulled from borosilicate glass (outer diameter, 1.5 mm; inner diameter, 0.87 mm; Hilgenberg) on a vertical puller (PC-10 model, Narishige) and filled with intracellular solution containing the following: 120 mM K-gluconate, 5 mM KCl, 10 mM HEPES, 4 mM Mg_2_ATP, 0.3 mM Na_4_GTP, 10 mM Na-phosphocreatine, pH 7.4, adjusted with KOH, and osmolarity of 275 mOsm. Dextran-labeled MNs were visualized using a fluorescence microscope (Axioskop FS Plus, Zeiss) equipped with IR-differential interference contrast optics and a CCD camera with frame grabber (Hamamatsu) and were then targeted specifically. Intracellular patch-clamp electrodes were advanced in the exposed portion of the spinal cord through the meninges using a motorized micromanipulator (Luigs & Neumann) while applying constant positive pressure. Intracellular signals were amplified with a MultiClamp 700B intracellular amplifier (Molecular Devices) and low-pass filtered at 10 kHz. In current-clamp recordings, no bias current was injected. Only motoneurons that had stable membrane potentials at or below -48 mV fired action potentials to suprathreshold depolarizations and showed minimal changes in series resistance (<5%) were included in this study. In some experiments neurons were passively filled with 0.25% neurobiotin tracer (Vector Labs) for post hoc analysis of their neurochemical identity. Spinal cords with neurobiotin-filled neurons were dissected out and transferred in 4% PFA solution at 4 °C for 3 h. The tissue was then washed extensively with PBS, and incubated in streptavidin conjugated to Alexa Fluor 555 (1:500; Invitrogen, SP-1120) and were used for detection of ChAT immunoreactivity following the protocol described above. The following drugs were added to the physiological solution: non-selective muscarinic receptor agonist muscarine (15 μM; Sigma, M104), m2-type selective muscarinic receptor antagonist methoctramine (10 μM; Sigma, M105) and m2-type preferential muscarinic receptor agonist oxotremorine-M (20 μM; Sigma, O100). All drugs were dissolved as stock solutions in distilled water. For all the electrophysiological recordings data analysis was performed using Spike2 (version 7, Cambridge Electronic Design) or Clampfit (Molecular Devices) software. The action potential voltage threshold of motoneurons was determined from the measured membrane potential at which the *dV/dt* exceeded 10 mV/msec.

#### *In vivo* swimming behavior

The swimming ability of zebrafish was tested using the open field test and the critical speed (U_crit_) test. U_crit_ is a measure of the highest sustainable swimming speed achievable by a fish. All zebrafish (n = 80) selected for the test displayed similar body length sizes and body weights. Animals were first anaesthetized in 0.03% tricaine methane sulfonate (MS-222, Sigma-Aldrich) in fish water and injected intraperitoneally (volume: 2 μl) with saline, muscarine (50μμ; 525 ng / g body weight) or/and methoctramine (40 μM; 2230 ng / g body weight; Figure S3). Treated animals were placed in the swim tunnel (5 L; Loligo systems, Denmark) to recover and acclimated at a low water flow speed (4.5 cm/sec) for 7 min. After, fish were given the U_crit_ test, subjecting the animals to time intervals of a certain flow velocity (increments of 4.5 cm/sec in 5 min steps) until the fish could not swim against the water current (fatigued; Figure S3). Fatigue was determined when fish stopped swimming and was forced against the rear net of the tunnel for more than 5 sec. Critical speed was then calculated using the following equation (Brett, 1964):

U_crit_ = U_fatigue_ + [U_step_ × (t_fatigue_ / t_step_)],

*Where:* U_fatigue_ = the highest flow velocity where fish swam the whole interval, Ustep = velocity increment, tfatigue = time elapsed at final velocity that fish swam in the last interval, tstep = time increment that is the duration of one interval. The critical speed was normalized to body length (BL) of the experimental animals and is given as BL/sec.

For the open field test, treated animals were placed in small dishes (diameter: 8 cm) and allowed to swim freely, while their swimming was recorded for 4 min. Analysis of 2 min swimming behavior was performed after optimization and implementation of wrMTrck, a freely available ImageJ plugin. The average velocity and maximum velocity was normalized to body length (BL) of the experimental animals and is given as BL/sec.

### Statistics

The significance of differences between the means in experimental groups and conditions was analyzed using the *One-way* ANOVA followed by *post hoc* Tukey test and the two-tailed Student’s *t*-test (paired or unpaired), using Prism (GraphPad Software Inc.) Differences were considered to be significant if p < 0.05. Data presented here are given as mean ± SEM.

## Acknowledgements

We are grateful to A. El Manira for providing the electrophysiology set-up and the zebrafish lines; and K. Meletis and A. Martin for critical assistance with *in situ* hybridization experiments. We also thank A. El Manira, C. Broberger, M. Carlén, G. Silberberg, and P. Williams for valuable discussion and comments of our manuscript. This work was supported by a grant from the Swedish Research Council (2015-03359 to K.A.), StratNeuro (to K.A.) Swedish Brain Foundation (FO2016-0007 to K.A.) and Längmanska kulturfonden (BA17-0390 to K.A.).

---

**Supplementary file 1.**
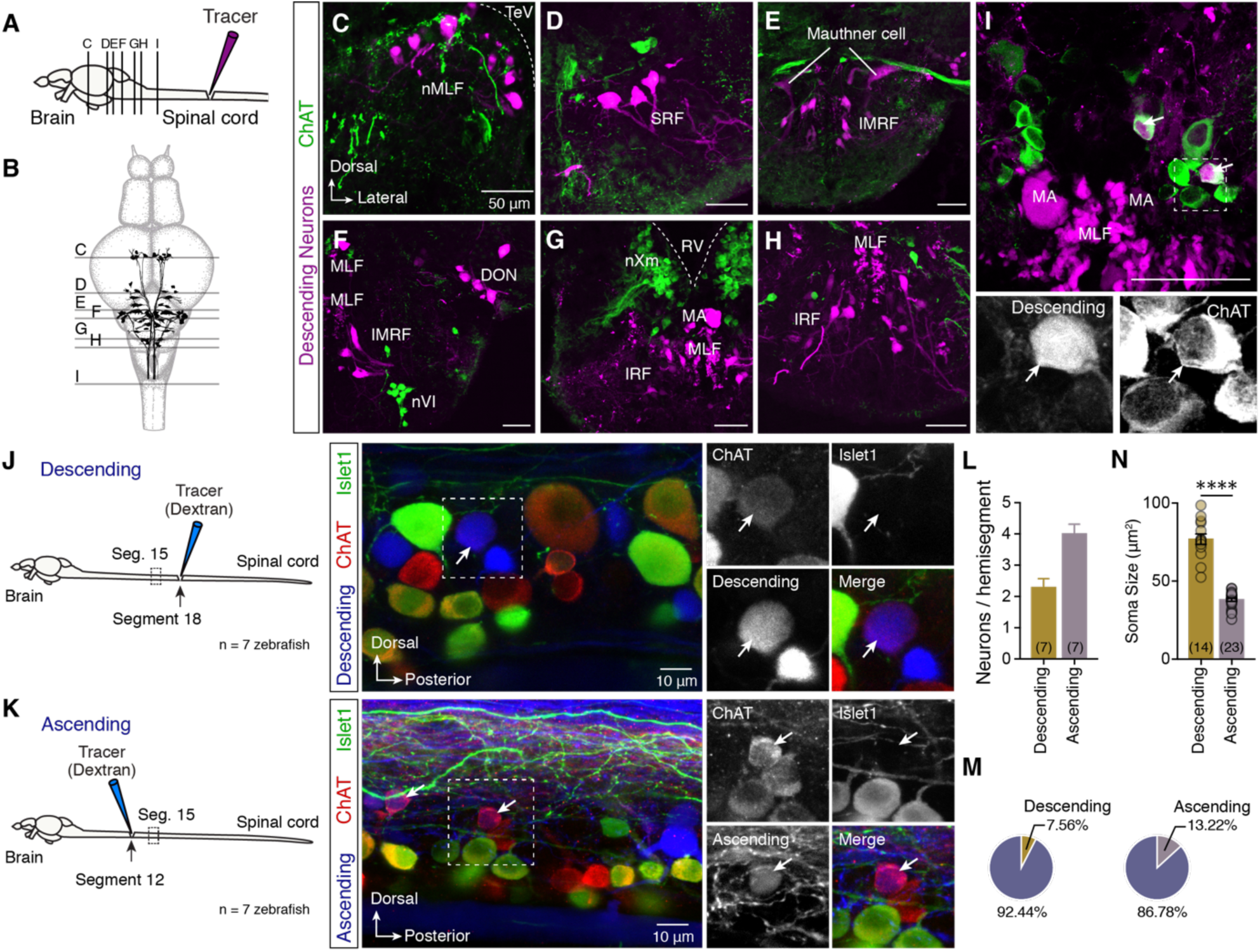
All the brain neurons descending to spinal cord are not cholinergic. (**A-B**) Injection of methylrhodamine dextran in the spinal cord retrogradely labels all the descending supra-spinal neurons (N = 8 zebrafish brains). Schematic representation of the distribution of adult zebrafish brain descending neurons with the level of sections that correspond to the following images in C-I. (**C-H**) Confocal images show that none of the descending labeled neuron is ChAT^+^ in all studied brain areas. (**I**) Few descending neurons (n = 4 neurons out of 8 brains) were found to be ChAT^+^ in the initial part of the spinal cord. Arrows indicate the double labeled neurons. (**J**) Injection of a retrograde dextran tracer in spinal segment 18 reveals the descending interneurons located in spinal cord segment 15. Few descending neurons that are cholinergic (ChAT^+^) and not motoneurons (Islet1^-^) were found in the spinal hemisegment of adult zebrafish. Arrow indicates cholinergic descending interneuron. (**K**) Injection of dextran retrograde tracer in spinal segment 12 reveals the ascending interneurons in spinal cord segment 15. Arrows indicate a small number of retrograde traced neurons that are cholinergic interneurons (ChAT^+^ Islet1^-^). (**L**) Analysis of the number of the cholinergic interneurons that possess a descending (2.28 ± 0.28 neurons / hemisegment; n = 7 zebrafish) or an ascending (4 ± 0.3 neurons / hemisegment; n = 7 zebrafish) axon. (**M**) Percentage of the descending and ascending cholinergic interneurons per hemisegment of spinal cord. (**N**) The descending and ascending cholinergic interneurons have non-overlapping soma sizes (t = 13.4, p < 0.0001, n = 37 neurons), suggesting that different populations of cholinergic interneurons are ascending or descending in adult zebrafish spinal cord. DON, descending octaval nucleus; IMRF, intermediate reticular formation; IRF, inferior reticular formation; MA, Mauthner axon; MLF, medial longitudinal fascicle; nMLF, nucleus of the medial longitudinal fascicle; nVI, abducens nucleus; nXm, vagal motor nucleus; RV, rhombencephalic ventricle; SRF, superior reticular formation; TeV, tectal ventricle. Data are presented as mean ± SEM; ****p < 0.0001.

**Supplementary file 2.**
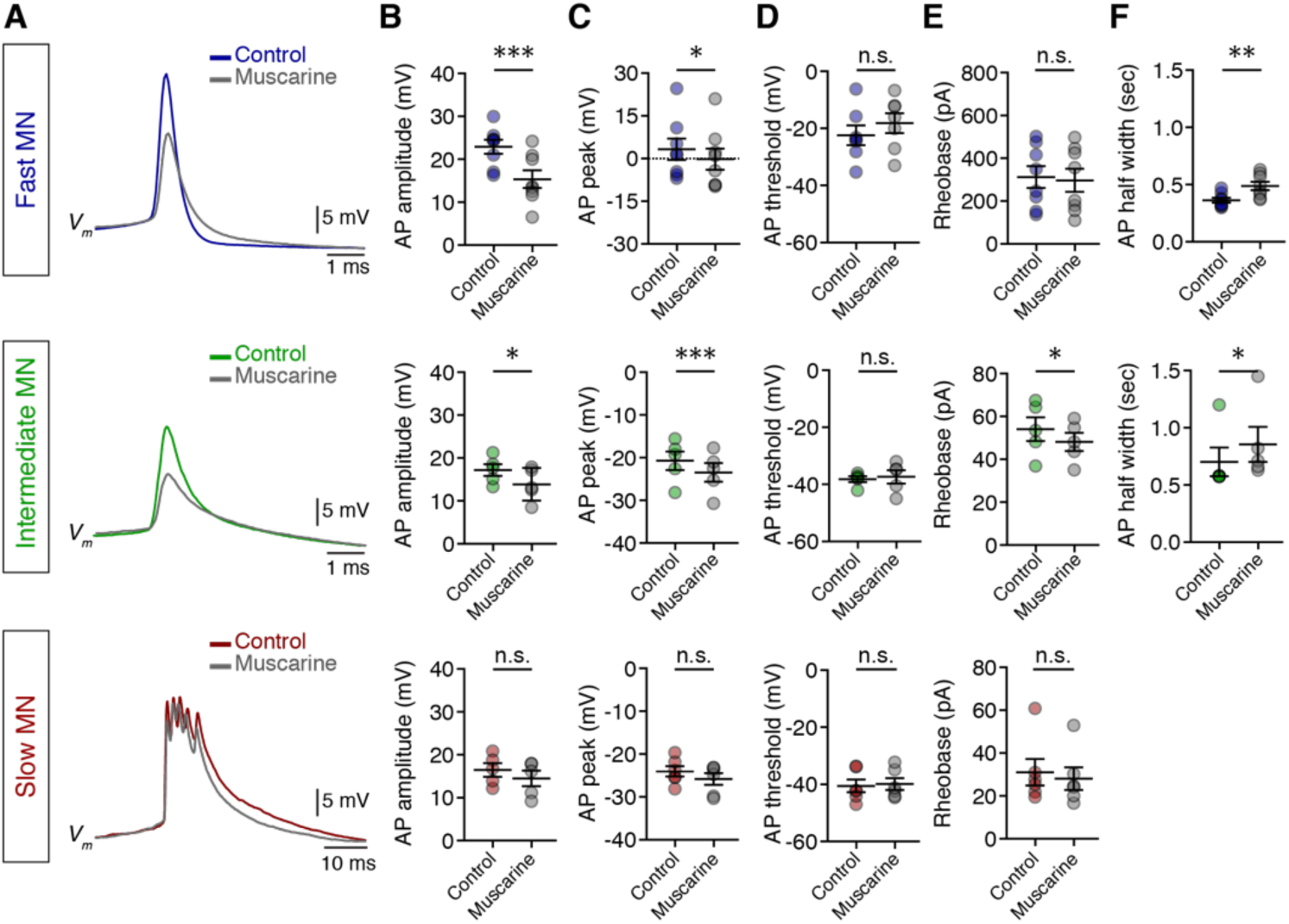
Changes in MN intrinsic biophysical properties after the bath application of muscarine. (**A**) Superimposed representative examples of the first action potential (AP) in fast and intermediate motoneurons and first burst of action potentials (APs) of slow motoneurons before (colored traces) and after (gray traces) muscarine. (**B**) The AP amplitude was significantly reduced by muscarine in intermediate (t = 4.33, p = 0.0123, n = 5) and fast motoneurons (t = 5.66, p = 0.0008, n = 8). (**C**) Following the reduction of AP amplitude also the AP peak was found to be hyperpolarized (Fast MNs: t = 2.44, p = 0.044, n = 8; intermediate MNs: t = 14.64, p = 0.0001). (**D**) Muscarine application was found not to affect the AP threshold. (**E**) Rheobase, the minimum current injection that generates an AP, was hyperpolarized in intermediate motoneurons (t = 4.24, p = 0.013, n = 5). (**F**) The half-width duration of the AP was significantly increased by muscarine in motoneurons firing single APs (intermediate MNs: t = 3.79, p = 0.019, n = 5; fast-MNs: t = 4.9, p = 0.0017, n = 8). Student’s paired t test. Data are presented as mean ± SEM; *p < 0.05; **p < 0.01; ***p < 0.001; n.s., non-significant.

**Supplementary file 3.**
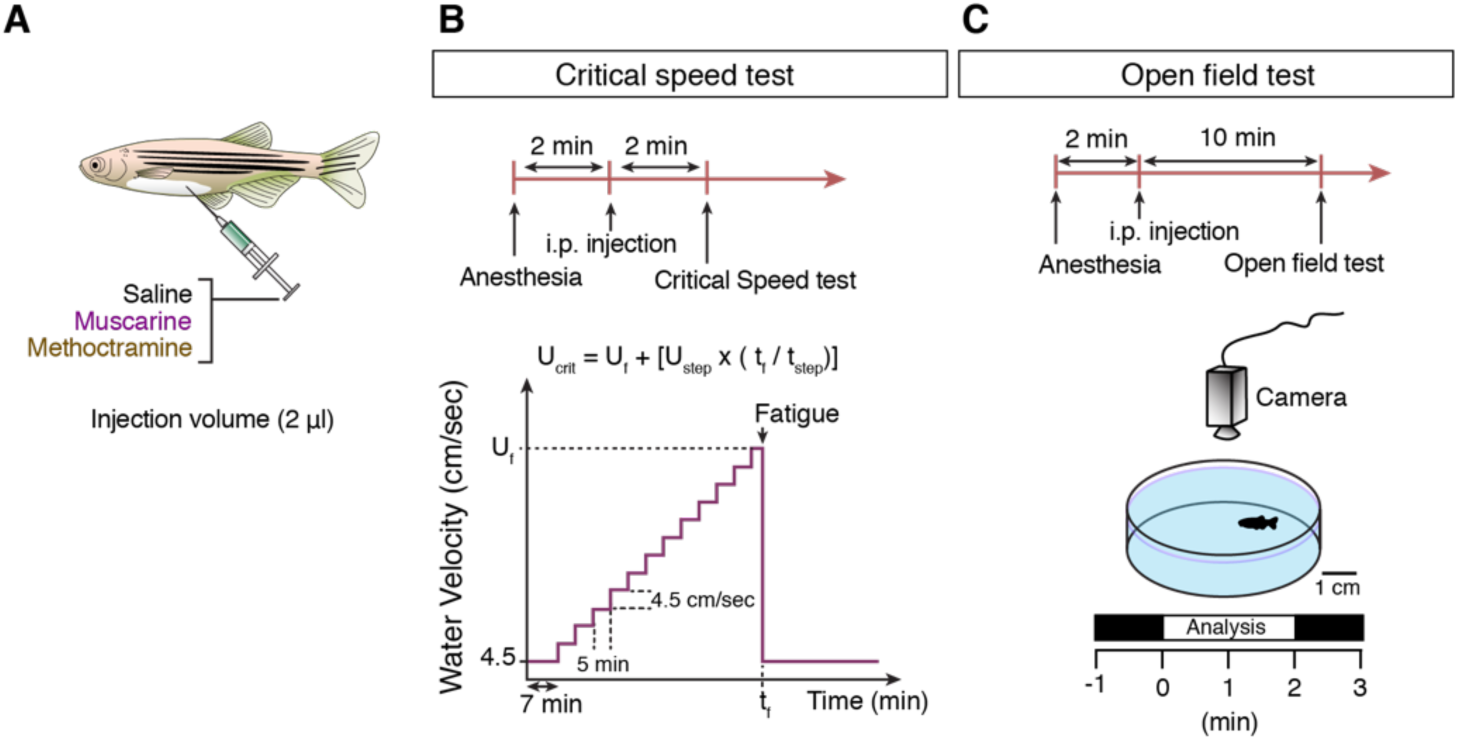
Experimental design to investigate the effect of mAChRs during *in vivo* locomotion. (**A**) Intraperitoneal administration of saline, muscarine and/or methoctramine in anesthetized adult zebrafish before the *in vivo* tests. (**B**) The critical speed test protocol. (**C**) Protocol for *in vivo* monitoring of adult zebrafish locomotor behavior.

